# Jumonji-C demethylase 2 interacts with a nucleosome component to modulate chromatin state in *Plasmodium falciparum*

**DOI:** 10.1101/2025.11.03.686342

**Authors:** Tatiane Macedo-Silva, Rosana Beatriz Duque Araujo, Luisa Fernanda Ortega Sepulveda, Juliane Cristina Ribeiro Fernandes, Gerhard Wunderlich

## Abstract

Epigenetic regulation plays a central role in the developmental control and antigenic variation of *Plasmodium falciparum*, yet the functions of many chromatin-modifying enzymes remain poorly understood. Here, we investigated the role of the putative Jumonji C (JmjC) domain–containing enzyme PfJmjC2 (PF3D7_0602800) in maintaining chromatin organization and histone modification balance during the asexual blood stages of the parasite. Using a conditional glmS ribozyme-mediated knockdown system, we achieved a ∼70% reduction in PfJmjC2 transcript and protein levels, which resulted in a marked delay in DNA replication and intraerythrocytic development. Immunoprecipitation assays revealed that PfJmjC2 physically interacts with histone H2A, suggesting an association with the nucleosome core. Upon PfJmjC2 depletion, the balance of epigenetic marks was disrupted, indicating a shift toward a more compact and transcriptionally repressed chromatin state. MNase–ChIP–PCR analysis further showed that PfJmjC2 associates with nucleosome-dense regions, including *var* gene promoters, although its reduction did not alter *var* gene transcription or switching patterns. Together, these findings indicate that PfJmjC2 contributes to chromatin organization and nucleosome stability rather than direct transcriptional control, highlighting its potential role in maintaining chromatin compaction and epigenetic homeostasis in *P. falciparum*.

## 1. Introduction

Malaria remains one of the most significant infectious diseases globally, caused by apicomplexan parasites of the genus *Plasmodium*. The most virulent species that infect humans, *P. falciparum*, continues to be responsible for the majority of severe cases and deaths. According to the World Health Organization (World malaria report 2024, n.d.), there were approximately 249 million malaria cases and around 608,000 deaths worldwide in 2023. Despite advances in prevention and treatment, progress has stalled in recent years due to emerging drug and insecticide resistance, disruptions from global health crises such as COVID-19, and environmental changes affecting mosquito transmission patterns. The parasite’s complex life cycle, alternating between mosquito and human hosts and spanning diverse cellular environments, demands tight regulation of gene expression at each developmental stage. In the human host, *P. falciparum* undergoes replication in hepatocytes and red blood cells (RBCs), with gene expression following a highly ordered, stage-specific transcriptional program (Bozdech et al., 2003; Le Roch et al., 2003). This transcriptional control is mediated by multiple factors, including plant-like Apetala2 (AP2) transcription factors (Balaji et al., 2005) and an array of chromatin-modifying enzymes such as histone acetyltransferases, deacetylases, methyltransferases, and demethylases (Coleman and Duraisingh, 2008; Croken et al., 2012). In *P. falciparum*, these chromatin modifiers play essential roles in regulating gene networks involved in differentiation, virulence, invasion, and nutrient transport (Crowley et al., 2011; Jiang et al., 2013; Mira-Martínez et al., 2013; Ukaegbu et al., 2014; Cortés and Deitsch, 2017; Duraisingh and Skillman, 2018). One of the best-studied examples of epigenetic regulation in P. falciparum involves the control of *var* genes, which encode the major virulence factor *P. falciparum* erythrocyte membrane protein 1 (PfEMP1). The expression of these genes is mutually exclusive and epigenetically regulated through heterochromatin marks, histone modifiers, and nuclear positioning (Tonkin et al., 2009; Cui and Miao, 2010; Ukaegbu et al., 2014). Previous genetic studies by Jiang and colleagues (Jiang et al., 2013) identified several histone-modifying enzymes that contribute to *var* gene transcriptional regulation. Among them, a putative Jumonji-like histone demethylase, PfJmjC2 (PF3D7_0602800), was disrupted without visible effects on parasite growth or *var* transcription levels. However, this study utilized phenotypically mixed parasite populations, limiting conclusions about *var* gene expression memory or dynamic chromatin regulation. Later, Zhang et al. (Zhang et al., 2018) reported that piggyBac insertional disruption of PfJmjC2 was tolerated, albeit with a measurable fitness cost. Furthermore, Matthews et al. (2021) demonstrated that Jumonji-domain-containing demethylases could be inhibited not only in asexual stages but also in gametocyte and gamete stages, underscoring their potential as therapeutic targets. Considering potential functional redundancy among *P. falciparum* chromatin modifiers—which may compensate for the loss of PfJmjC2 in knockout lines—we aimed to characterize the immediate consequences of temporary PfJmjC2 depletion. Here, we generated a transgenic parasite line in which the endogenous PfJmjC2 open reading frame was tagged at its 3′ terminus with an HA epitope and a glmS ribozyme sequence (Prommana et al., 2013). Upon glucosamine treatment, glmS activation induced rapid, though incomplete, degradation of the PfJmjC2 transcript. This system enabled us to investigate the short-term molecular effects of PfJmjC2 depletion on histone modification patterns, chromatin organization, and gene expression in *P. falciparum*.

## 2. Results

### 2.1. C-terminally tagged PfJmjC2 is stably expressed throughout the blood-stage cycle and localizes to the nucleus

To examine the stage-specific expression of PfJmjC2, we generated clonal parasite lines carrying C-terminal HA or GFP-HA and HA-glms tags at the endogenous *PfJmjC2* locus (Figure S1). Synchronized transfectant parasite lines were obtained through successive plasmagel and sorbitol treatments. Relative PfJmjC2 transcript levels were quantified by reverse transcription-qPCR (RT-qPCR), showing consistent expression across ring, trophozoite, and schizont stages (Figure 1A). Western blot analysis using an anti-HA antibody confirmed that PfJmjC2 protein is also present in all tested intraerythrocytic forms (figure 1B), consistent with available proteomic and transcriptomic data from PlasmoDB (www.plasmodb.org). Fluorescence microscopy of parasites expressing a GFP-tagged version of PfJmjC2 revealed a strong nuclear signal, confirming that the protein localizes to the nucleus (figure 1C)x’. This nuclear localization pattern demonstrates that the addition of C-terminal tags does not interfere with proper targeting of PfJmjC2. These findings establish that PfJmjC2 is stably expressed throughout the asexual blood-stage cycle and functions as a nuclear, probably chromatin-associated protein in *P. falciparum*.

**Figure 1.**
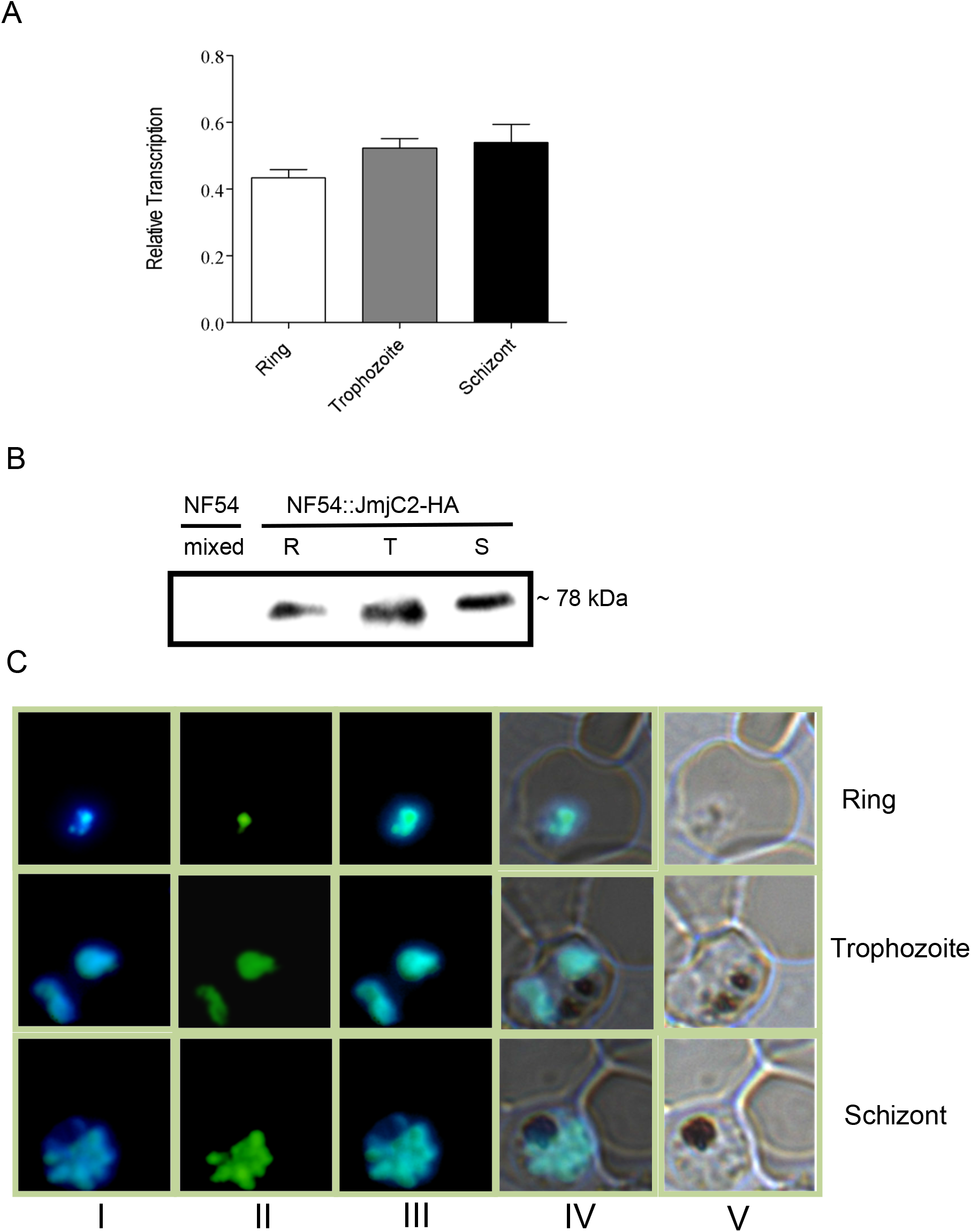
PfJmjC2 is expressed throughout the intraerythrocytic cycle and localizes to the nucleus. **A:** Quantitative real-time PCR showing relative PfJmjC2 transcript levels at ring (white, 4–10 hpi), trophozoite (grey, 18–26 hpi), and schizont (black, 34–42 hpi) stages. Transcript levels were normalized to *seryl-tRNA ligase* (PF3D7_0717700). Parasites were synchronized twice prior to RNA collection. **B:** Western blot of NF54::PfJmjC2-HA-glmS parasites showing stage-specific expression in ring (R), trophozoite (T), and schizont (S) stages, detected with anti-HA. NF54 parasites from asynchronous culture were used as a control. **C:** Fluorescence microscopy of NF54::PfJmjC2-HA-GFP-glmS parasites at ring, trophozoite, and schizont stages. Panels: (I) DAPI-stained nuclei, (II) GFP-tagged PfJmjC2, (III) overlay of I and II, (IV) overlay of III and V, and (V) bright field.

### 2.2. PfJmjC2 interacts with histone H2A and modulates histone modification patterns

Previous studies have shown that JmjC domain–containing proteins can demethylate several histone lysine residues, including H3K9me3 and H3K4me1/2/3, and may also interact with histone H2A (Markolovic et al., 2016; Matthews et al., 2021; Cui et al., 2008). To determine whether the plasmodial PfJmjC2 associates with H2A, we performed immunoprecipitation of PfJmjC2–HA using anti-HA antibodies, followed by detection with an anti-H2A antibody. As shown in Figure 2A, a protein band corresponding to the molecular weight of histones was detected in the immunoprecipitate, confirming that H2A co-precipitates with PfJmjC2. This interaction positions PfJmjC2 close to the nucleosome core, suggesting a role in regulating nucleosome stability and histone turnover. To assess whether PfJmjC2 depletion affects histone modification patterns, we examined the abundance of the marks H3K4me1, H3K9me3, and H3K9ac in total parasite extracts after glucosamine-induced knockdown of PfJmjC2–HA. Histone modifications were analyzed by Western blot using specific antibodies, and their relative intensities were normalized to total H3 (Figure 2B). Densitometric analysis suggested shifts in these marks (Figure 2C). An increase in the silencing mark H3K9me3, which is associated with virulence gene regulation and parasite developmental control, was observed, consistent with a loss of demethylase activity. Concomitantly, H3K4me1, an activation mark enriched at regulatory enhancer regions, and H3K9ac, typically found at the 5′ regions of active genes, both decreased, indicating reduced chromatin accessibility and transcriptional repression upon PfJmjC2 depletion. Together, these findings demonstrate that PfJmjC2 physically associates with histone H2A and plays a central role in maintaining histone modification balance. These results suggest that PfJmjC2 contributes to chromatin organization and histone mark homeostasis in *P. falciparum*, although its specific molecular targets and regulatory outcomes may vary depending on the chromatin region or developmental stage of the parasite.

**Figure 2.**
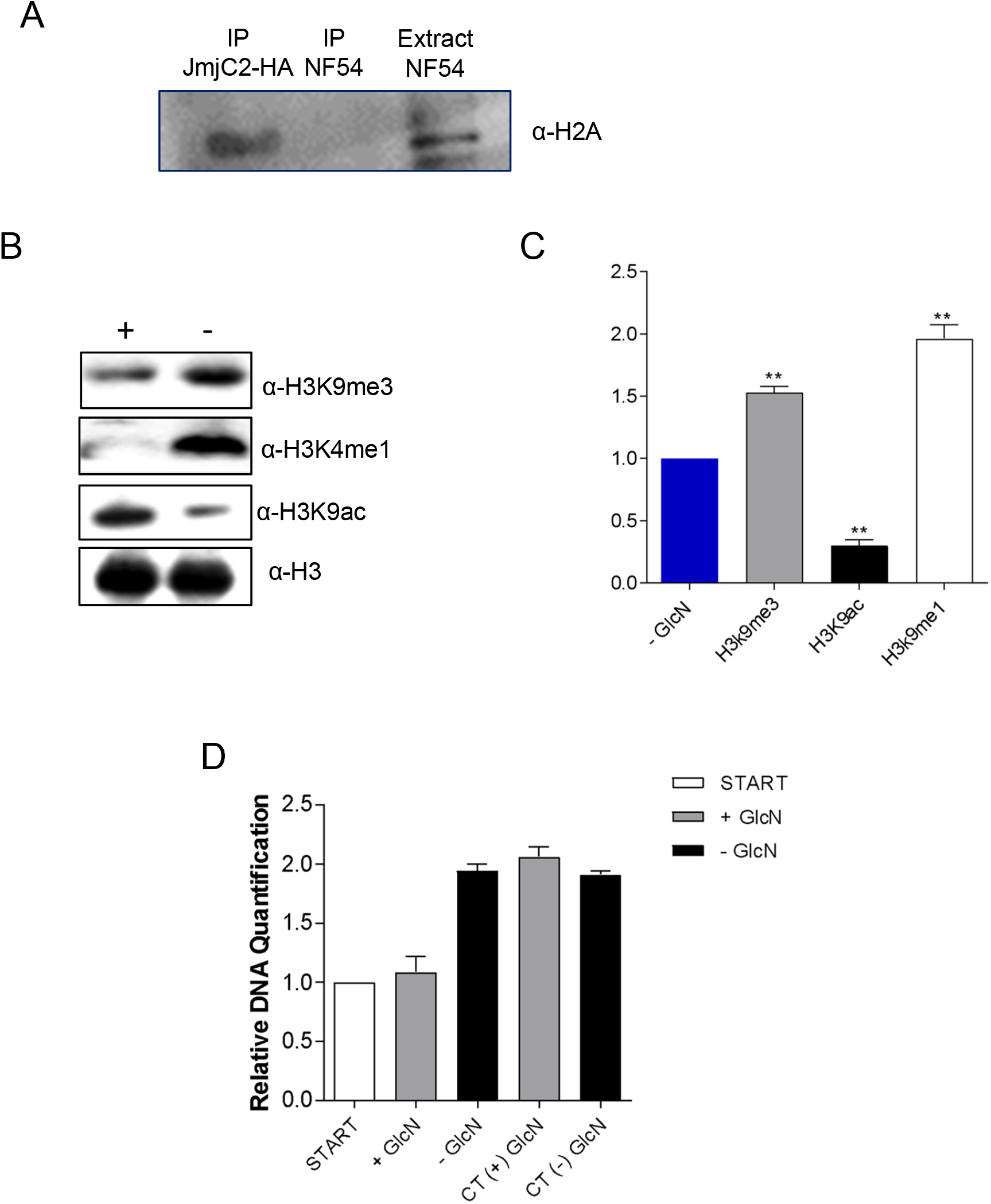
PfJmjC2 can be knocked down without causing growth defects. **A:** Western blot of NF54::PfJmjC2-HA-glmS and NF54 trophozoite-stage parasites cultured ± 2.5 mM GlcN. Protein extracts were probed with anti-HA (left panel, ∼70 kDa) and anti-ACT (right panel, ∼40 kDa). **B:** Densitometric analysis of HA signals normalized to ACT using ImageJ. **C:** Quantification of PfJmjC2 transcripts after RNA destabilization upon GlcN treatment. Trophozoites (30 hpi) were analyzed after 24 h of exposure to 2.5 mM GlcN, initiated at the ring stage (6 hpi). **D:** Growth curves of NF54::PfJmjC2-HA-glmS and wild-type NF54 parasites treated with various GlcN concentrations and analogs, showing growth inhibition only at the highest concentrations in NF54::PfJmjC2-HA-glmS parasites.

### 2.3. Conditional knockdown of PfJmjC2 reduces its expression and delays parasite development

To elucidate the function of PfJmjC2, we generated a conditional knockdown line and treated tightly synchronized ring-stage parasites 5mM glucosamine (GlcN) for 24 hours. This treatment resulted in a 70–75% reduction in PfJmjC2 transcript and protein levels, as confirmed by RT-qPCR and Western blot analyses, respectively (Figure 3 A/B/C). Importantly, this concentration of GlcN did not affect the growth or morphology of wild-type parasites, indicating that the observed effects were specific to PfJmjC2 depletion (Figure 3). Parasites with reduced PfJmjC2 expression exhibited a pronounced developmental delay, remaining in pre-replicative stages while control parasites had already initiated DNA replication after 24h of treatment (Figure 2D). Together, these findings demonstrate that PfJmjC2 is required for proper intraerythrocytic development and suggest a critical role in facilitating timely progression through the S phase.

**Figure 3.**
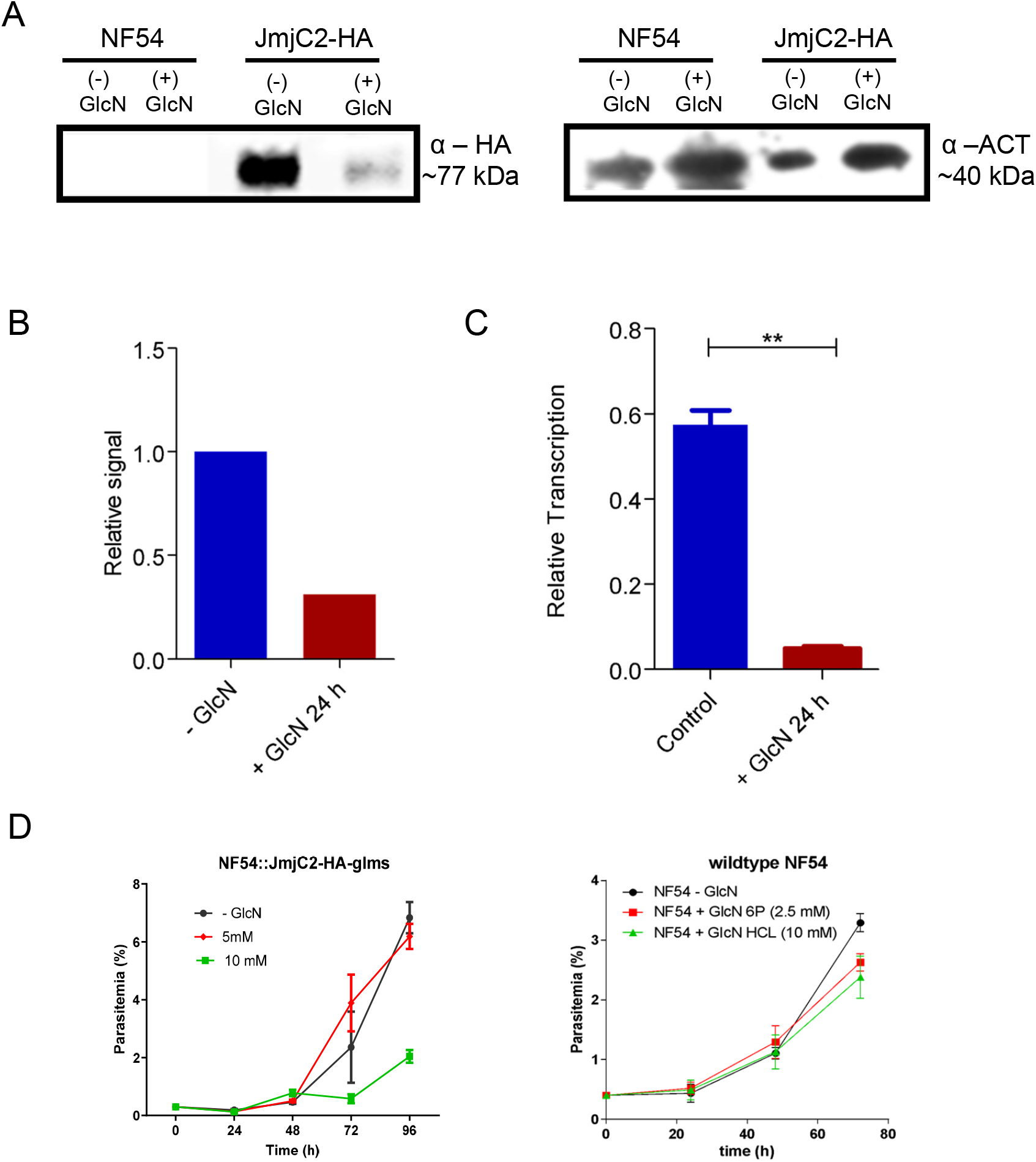
PfJmjC2 associates with histone H2A and influences global histone modifications. **A:** Western blot showing that H2A co-immunoprecipitates with PfJmjC2-HA. IP PfJmjC2-HA corresponds to anti-HA–immunoprecipitated material, while IP NF54 represents wild-type parasites processed with anti-HA. The detected H2A protein (∼17 kDa, arrow) confirms its presence in complex with PfJmjC2-HA. **B:** Western blot showing the effect of PfJmjC2 transcript depletion on global H3K9me3 and H3K9ac levels in NF54::PfJmjC2-HA-glmS parasites ± GlcN treatment, using anti-H3 as a loading control. **C:** Densitometric quantification of H3K9me3 and H3K9ac levels after 24 h ± GlcN treatment. Values represent the mean of three independent experiments; differences were statistically significant at *p* < 0.05 (*) or *p* < 0.01 (**), Student’s *t*-test. **D:** Relative DNA quantification by real-time PCR showing that, after 24 h of GlcN treatment, wild-type parasites continue DNA replication, whereas NF54::PfJmjC2-HA-glmS parasites do not (*n* = 3).

### 2.4. PfJmjC2 associates with nucleosome-dense chromatin and var gene loci

DNA obtained from PfJmjC2–HA MNase-ChIP (Figure 4A) assays was used for PCR amplification with primers targeting *var* gene promoter regions. A strong PCR signal corresponding to a DNA fragment of approximately 300 bp was detected from the MNase-digested chromatin (Figure 4B). This fragment size is consistent with dinucleosomal protection, indicating that *var* promoters remain shielded from nuclease digestion and are embedded within compact chromatin regions. Such protection suggests that these promoter regions are nucleosome-dense and not part of open, transcriptionally active chromatin domains. These findings support a model in which PfJmjC2 preferentially associates with highly compact chromatin, including *var* gene promoters, where it may contribute to maintaining the repressive chromatin configuration important for *var* silencing.

**Figure 4.**
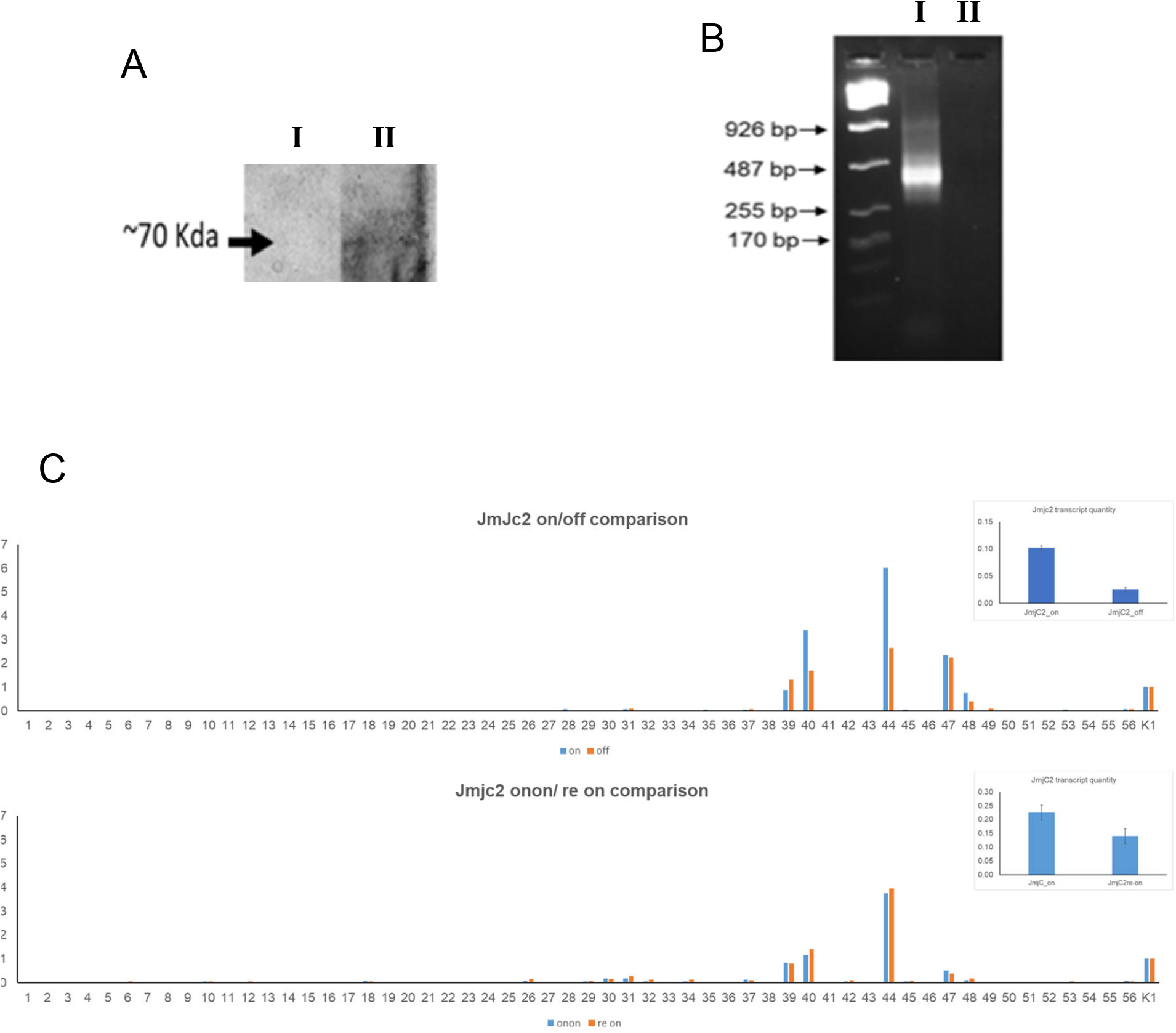
var gene promoters are nucleosome-enriched, and var transcription is largely unaffected by partial PfJmjC2 reduction, maintaining transcriptional memory. **A:** Western blot showing successful immunoprecipitation of PfJmjC2-HA using an anti-HA antibody, with a specific band detected at approximately 70 kDa in lane II (I: input; II: immunoprecipitated fraction). **B:** MNase-PCR analysis confirming that *var* promoter regions remain associated with nucleosome-dense chromatin under PfJmjC2 pull-down conditions. Lane I represent DNA recovered from the PfJmjC2 immunoprecipitation, and lane II corresponds to input DNA, both used as amplification templates. The positive band in lane 1 indicates successful PCR amplification, while lane 2 shows no amplification under the same conditions. **C:** Quantitative real-time PCR showing *var* gene expression profiles in parasites with reduced PfJmjC2 levels, in controls, and after restoration of normal PfJmjC2 expression, with each bar representing a distinct *var* locus. Gene identifiers are listed in Supplementary Table S1. These findings indicate that *var* promoters are enriched in nucleosomes and that partial PfJmjC2 depletion does not significantly alter *var* transcription, supporting the maintenance of transcriptional memory despite reduced PfJmjC2 activity.

### 2.5. Despite its association with *var* gene promoters decreased levels of PfJmjC2 do not alter *var* transcription patterns

Considering PfJmjC2’s influence on H3K9 modification and its association with chromatin regions containing *var* genes, we examined whether *var* transcription was affected by PfJmjC2 knockdown. NF54::PfJmjC2-HA-glmS parasites were selected for cytoadherence to human CD36-expressing CHO cells, which reproducibly induces transcription of specific *var* genes, mainly of the centromeric upsC type. Selected parasites were then treated or not with glucosamine and harvested at ring stage (4–10 h post-invasion). RT-qPCR analyses revealed no major changes in the overall *var* transcription pattern following PfJmjC2 knockdown (Figure 4C and Figure S2). The same dominant *var* transcripts persisted in both conditions, with only a modest reduction in transcript abundance for the most highly expressed *var* genes such as the centromeric loci on chromosome 4 (PF3D7_1240400 and PF3D7_0412400) (Figure 4C). Extended glucosamine exposure (10 mM for 96 h) followed by recovery also did not induce detectable *var* gene switching, and this result was consistent even after >15 replication cycles post-panning over CHO-CD36 cells. These findings demonstrate that although PfJmjC2 associates with subtelomeric *var* loci, its depletion does not alter *var* promoter activation or switching activity. The slight reduction in *var* transcript abundance likely reflects global chromatin condensation and reduced transcriptional accessibility rather than direct *var* transcription regulation. Therefore, PfJmjC2 seems to function as a nucleosome-level chromatin regulator that may support genome-wide transcriptional balance but is not directly responsible for *var* gene expression or epigenetic memory maintenance.

## 3. Discussion

To date, three Jumonji C (JmjC)-domain histone demethylases have been identified in the *P. falciparum* genome. These enzymes represent promising epigenetic drug targets, as their inhibition leads to parasite death in both asexual and sexual stages (Matthews et al., 2021). In this study, we characterized the immediate consequences of PfJmjC2 depletion on chromatin organization, histone modification balance, and transcriptional control, with a particular focus on variant surface antigen (*var*) genes. Previous studies (Jiang et al., 2013; Zhang et al., 2018) suggested that PfJmjC2 is dispensable for parasite survival, although its absence imposes a mild fitness cost. Using a conditional glmS ribozyme-mediated knockdown system, we found that PfJmjC2 depletion led to a reproducible developmental delay of approximately 12–16 hours, coinciding with postponed DNA replication. This phenotype suggests that PfJmjC2 contributes to the chromatin relaxation required for timely replication and transcription. Supporting this idea, PfJmjC2 knockdown altered the equilibrium of histone marks, increasing repressive modifications (H3K9me3) while reducing the activating mark H3K9ac. These coordinated shifts suggest a role in maintaining chromatin fluidity and nucleosome plasticity essential for transcriptional accessibility. Immunoprecipitation assays further revealed that PfJmjC2 physically interacts with histone H2A, implicating a role at the nucleosome core. This association likely supports nucleosome stability and turnover, facilitating the dynamic chromatin environment necessary for replication and gene expression. Comparable interactions between JmjC-domain proteins and H2A have been described in yeast (Huang et al., 2015), suggesting a conserved mechanism across eukaryotes. Given that the H2A variant H2A.Z is a hallmark of active chromatin in *P. falciparum* (Bartfai et al., 2010; Hoeijmakers et al., 2013), our data point to a stabilizing role of PfJmjC2 in nucleosome organization and chromatin balance throughout the parasite life cycle. Despite PfJmjC2 enrichment at several *var* promoters within subtelomeric chromatin, its reduction did not alter *var* gene switching or the dominant expression pattern. The persistence of the same *var* transcripts in both control and PfJmjC2-reduced parasites, with only minor transcript variation, indicates that PfJmjC2 influences chromatin structure globally but does not directly regulate *var* promoter activation. Consistent with this, MNase-PCR analysis demonstrated that *var* promoter regions remain associated with nucleosome-dense chromatin. These findings suggest that transcriptional memory at *var* loci is maintained independently of PfJmjC2 activity within nucleosomes. Since *var* gene expression is largely governed by PfHP1-mediated heterochromatin architecture, this process is likely more influenced by JmjC1 (Gockel et al., 2025). Thus, our data indicate that PfJmjC2 acts at the nucleosome level, possibly contributing to chromatin flexibility rather than the establishment of heterochromatin boundaries. This study provides new mechanistic insight into PfJmjC2 function using a conditional, short-term depletion system that avoids compensatory adaptations observed in long-term knockout lines. The integration of multiple assays—transcriptomics, histone mark profiling, MNase-PCR, and protein interaction— indicated chromatin composition changes with functional outcomes in replication and transcription. Nonetheless, limitations remain. PfJmjC2 depletion was partial (∼70%), and residual enzyme activity may have buffered the emergence of stronger phenotypes. Future studies employing a more complete knockdown or knockout strategy will be important to fully elucidate the impact of PfJmjC2 loss on parasite growth and gene regulation. Moreover, single-cell and time-resolved chromatin profiling approaches (e.g., scRNA-seq or ATAC-seq) could uncover transient or heterogeneous responses that are obscured in bulk analyses. Finally, the enzymatic specificity of PfJmjC2 toward individual histone residues remains to be biochemically validated. Together, our findings suggest that PfJmjC2 is a chromatin-associated enzyme that interacts with histone H2A and likely functions to preserve histone modification balance and nucleosome stability. Its reduction leads to delayed DNA replication and alterations in histone modification levels, indicating that PfJmjC2 may play a role in the formation or maintenance of dense, compact chromatin. These results refine and extend the conclusions of Gockel et al. (Gockel et al., 2025), demonstrating that PfJmjC2 acts at the nucleosome core level to sustain chromatin flexibility and transcriptional homeostasis, complementing PfJmjC1 role in higher-order heterochromatin organization. Despite remaining questions, this study establishes PfJmjC2 as a regulator of chromatin dynamics and underscores the importance of nucleosome-level mechanisms in malaria epigenetic regulation.

## 4. Material and Methods

### 4.1. Parasite culture and synchronization

*P. falciparum* lineage NF54, provided by Mats Wahlgren (Karolinska Institutet, Sweden), was used throughout the experiments. Blood stage parasites were maintained in RPMI supplemented with 0,23% NaHCO3, 0.5% Albumax 1 (Gibco, Rockville MD), and human B+ erythrocytes in a defined gas mixture (90% N2, 5% O2 and 5% CO2 (Trager and Jensen, 1976). The synchronization of parasites was done by plasmagel floatation (Lelièvre et al., 2005) of mature trophozoites followed by sorbitol lysis (Lambros and Vanderberg, 1979) of ring-stage parasites after reinvasion. Before each plasmagel floating or cytoadherence selection assay, parasites were cultivated for at least 48 h in the presence of 5% inactivated human B+ plasma and 0.25% Albumax 1. Usage of human blood and plasma was granted by the local Ethics Committee for Experiments involving humans (CEPSH) at the Institute for Biomedical Sciences at the University of São Paulo (process No. 874/2017).

### 4.2. Plasmid constructs and transfection

The 3’ end of the putative JumonjiC2 encoding gene (PlasmoDB ID PF3D7 _0602800) was PCR-amplified using the following oligonucleotides: forward AGATCTATATGGTTACATTATGATATACC, reverse CTGCAGGTATATAATCTTCATCTAGGTTT (introduced BglII and PstI restriction sites underlined). The amplicons were cloned in pGEM T easy vectors (Promega Inc.) and sequenced. Then, the corresponding JmjC2-3’-encoding fragment was excised using BglII and PstI (Thermo Fisher Scientific) and transferred to pTEXHA-glmS or pTEX-GFP-HA-glmS vectors (de Koning-Ward et al., 2009) digested with the same enzymes. Recombinant plasmids pJmjC2-HA-glms or pJmjC2-GFP-HA-glmS were grown to high quantities using the Maxiprep protocol (Sambrook, J., 2012) and used for transfections. Transfection of parasites was done using the protocol described by Hasenkamp and colleagues (Hasenkamp et al., 2012), with the exception that 1-2x107 parasites were used for reinvasion in plasmid-loaded RBC. Transfected parasites were grown using 2.5 nM WR99210 (a gift from Jacobus Inc., USA). For the integration via single crossover recombination, transfected parasite lines were cultivated for 14-20 days without WR99210, after which the drug was added again. Normally, after three cycles locus-integrated parasite lines were obtained. These were cloned by standard limiting dilution (Rosario, 1981) using FACS quantitation of parasites (Macedo-Silva et al., 2021).

### 4.3 Southern Blot Analysis

Genomic DNA was isolated from the WT (NF54) and NF54::JmjC2-HA-glmS parasites, using the Wizard® Genomic DNA Purification Kit (Promega). 5 μg of each gDNA and 15 ng of plasmid DNA (pDNA) from the pJmjC2-HA-glmS construct were digested using Bglll. The probe used the fragment containing the glmS sequence, which was amplified from the pJmjC2-HA-glmS plasmid using standard PCR conditions with digoxigenin-dUTP (DIG High Prime DNA Labeling and Detection Starter Kit I (Roche Diagnostics). The oligonucleotide primers used were 5′-ctcgagTAATTATAGCGCCCGAACTAAGC-3′ and 5′-ggtaccAGATCATGTGATTTCTCTTTG-3′, respectively. The Southern procedure was performed following the protocol provided by the manufacturer of the labeling kit (Roche), using Hybond N membranes (Amersham/GE Healthcare) and a hybridization temperature of 44 °C. Hybridized DIG-labelled DNA fragments were then detected by an antiDIG-alcaline Phosphatase antibody and detected by chemiluminescence, exposing X-ray films (GE Healthcare) for 15-30 minutes (Supplementary Figures).

### 4.4 Total RNA preparation

Tightly Plasmagel/Sorbitol synchronized parasites were harvested in ring, trophozoite, or schizont forms. IRBC were treated with 0.1% Saponin for 10 min at RT and then pelleted at 5000 g/4 °C for 10 min, washed once in 1 ml PBS and then resuspended in a final volume of 100 µl TE. Afterward, 1 ml Trizol (Invitrogen) was added and the sample was vortexed for 30 s and stored at −80 °C until use. Total RNA was prepared following the Trizol protocol provided by the manufacturer (Invitrogen). The final total RNA was dissolved in 20 µl RNAse free water, and RNA quality was checked by TBE-gel electrophoresis and its concentration was measured using a Nanodrop device (Thermo Scientific, USA) and stored at −80 °C until use.

### 4.5 Real-time PCR with parasite-derived cDNA

For transcript quantification, oligo pairs corresponding to the NF54 JmjC2 gene available in plasmoDB were used: forward GAAAAACTTTGATGAGTACGCAAA and reverse TCGTATGTCCTGGCTGTTTTTA. These oligonucleotides were designed using the Primer3 web application using settings identical to those used for the design of the endogenous control transcript oligos. For this, settings were set to a hybridization temperature of 58-62°C, oligos length of 22 nt, amplicon length of 80-120 nt, and 30-70% GC content, leaving the further settings at default values. Total RNA was converted to cDNA for a final concentration of 50 ng/µl, RNAs were then treated with DNAse1 (Thermo-Fermentas) and cDNA synthesis was done using reverse transcriptase RevertAid (Thermo-Fermentas) using random oligos as published earlier. The RT-qPCR amplification was done using 5xHotfire Pol SYBR Mix or EVA green 5xFirePol Mix (Solis Biodyne Inc.) on a Quant Studio 3 thermocycler (Thermo Scientific). Relative transcript quantities were then calculated by the 2−ΔCt method (Livak and Schmittgen, 2001) using the endogenous control seryl-tRNA ligase transcript (PlasmoDB PF3D7_0717700, herein termed “K1”). Sequences of real time PCR var oligos were the same as developed by Salanti and colleagues (Salanti et al., 2003) (Supplementary Table S1).

### 4.6 Knockdown assay for glmS-mediated transcript destabilization and phenotype observation

Glucosamine/HCl (Sigma-Merck) diluted in sterile water was added at the given final concentrations and parasites were incubated under normal growth conditions for two reinvasion cycles. Experiments were done in biological duplicates or triplicates.

### 4.7 Immunoblotting

For the detection of recombinant proteins in transgenic parasite lines, whole parasite protein extracts were prepared from saponin-lysed IRBCs as described in Methods in Malaria Research (Ljungström et al., 2008). Proteins were loaded on standard discontinuous SDS-polyacrylamide gels and transferred to Hybond C membranes (GE Healthcare/Amersham). After blocking with 4% skimmed milk in 1xPBS/0.1% Tween20, HA-tagged proteins were recognized using a murine antiHA antibody (Sigma-Merck) followed by an antiMouse IgG-peroxydase antibody (KPL or Sigma-Merck). Blots were exhaustively washed with PBS/Tween between incubations and finally incubated with ECL substrate (GE Healthcare). As a loading control, a murine anti-Pf-Aspartyl-Carbamoyl-transferase (ACTase) antibody (in-house produced mouse antiserum) was used. Chemoluminescent signals were either captured in an ImageQuant (GE) apparatus or on Amersham/GE X-ray films and intensities were quantified using ImageJ software (NIH). The obtained values were normalized using the signals generated by ACTase.

### 4.8 Fluorescence and immunofluorescence microscopy

To visualize green fluorescent protein or light emitted by fluorescent dyes, 100 µl of parasite cultures were centrifuged and washed 1x with PBS and resuspended in 100 µl PBS and incubated with DAPI (Sigma-Merck, 10 µg/ml) for 20 min. Immediately before analysis, the cells were transferred to slides and covered by coverslips. Image analysis was performed on a Zeiss Axio Observer ZV1 microscope. For immunofluorescence, cells were fixed in 4% paraformaldehyde and 0.0075% glutaraldehyde, followed by exposure to 125 mM glycine pH 7, 0.05% TX-100 in PBS for permeabilization. Primary antibodies were added at 1:100 in 3% BSA for 3 h and goat anti-mouse secondary antibodies (Molecular Probes, AlexaFluor 488 and 594) at 1:2000 for 1 h. Imaging was then performed as described above.

### 4.9 Immunoprecipitation of JmjC2-HA

Whole parasite protein extracts were prepared from saponin-lysed IRBCs as described in Methods in Malaria Research (Ljungström et al., 2008). Subsequently, IP of HA-tagged targets and associated proteins was performed using the µMACS HA Isolation Kit following the manufacturer’s instructions (Miltenyi Biotech). The samples of anti-HA-immunoprecipitated proteins were loaded on standard discontinuous SDS-polyacrylamide gels 12% and detected by anti-H2A antibodies (Abcam, ref No. 88770). Immunoprecipitated proteins were detected in gels by the silver-stain method.

### 4.10 MNase-Chip-PCR

Trophozoite-enriched cultures of NF54::PfJmjC2-HA-glmS parasites were washed three times with PBS and incubated for 10 minutes with 0.1% saponin. Crosslinking was performed with 1% formaldehyde for 15 minutes at 37 °C with agitation, then quenched with 0.125 M glycine for 5 minutes on ice. Parasites were pelleted by centrifugation at 4,000 rpm for 15 minutes at 4 °C and washed three times with PBS. Chromatin digestion was performed using 2 U/mL micrococcal nuclease (MNase) for 1 hour at 37 °C. Protein immunoprecipitation was then carried out with the MultiMACS HA Isolation Kit (12×8) according to the manufacturer’s instructions. Briefly, parasite pellets were resuspended in lysis buffer and incubated with anti-HA monoclonal antibody conjugated to µMACS MicroBeads for 30 minutes on ice. Samples were subsequently placed in the µMACS™ Separator for 30 minutes, washed four times, and eluted to recover the immunoprecipitated protein/DNA complexes. Input samples used as procedural controls were withdrawn prior to immunoprecipitation and processed identically, except that no antibody was added. DNA was extracted overnight at 65 °C in an extraction buffer (0.1 M EDTA pH 8, 1% SDS, and 200 µg/mL proteinase K), followed by phenol–chloroform purification. Purified nucleosomal DNA was then used as a template for PCR with primers corresponding to *var* gene promoter regions.

## 5. Acknowledgements

Research in GW’s lab is funded by research grants from FAPESP (grants 2015/17174-7, 2017/24267-7 and 2021/13727-2). GW and JCS are CNPq research fellows. TMS, RBDA, and JCRF were supported by CNPq fellowships. LFOS is supported by a doctoral fellowship from CAPES.

## 6. Authors contributions

TMS, JCRF, and RBDA created and characterized parasite lines, conducted part of the real-time experiments, blots and microscopy, prepared ChIPseq and RNAseq samples and proteomic analysis, APB, LFOS, and GW conducted real-time *var* transcript analysis, Conceptualization: TMS, RBDA, and GW. GW analyzed RNAseq results. The authors thank the Facilities Center for Research (CEFAP) at ICB-USP for generating sequencing results. TMS and GW drafted the manuscript.

## Data availability

Generated Fastq files are available upon request.

