## Supplemental figure 1 and 2 and table 1 for "Jumonji-C demethylase 2 interacts with a nucleosome component to modulate chromatin state in *Plasmodium falciparum*"

### Supplementary Figure S1

A

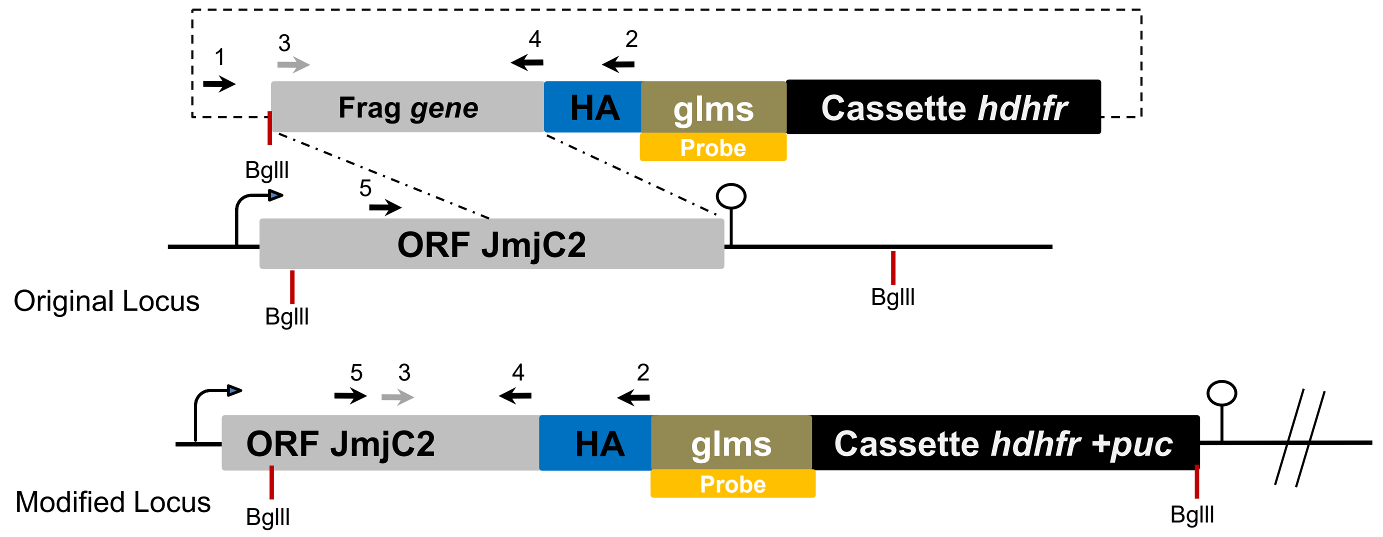


B

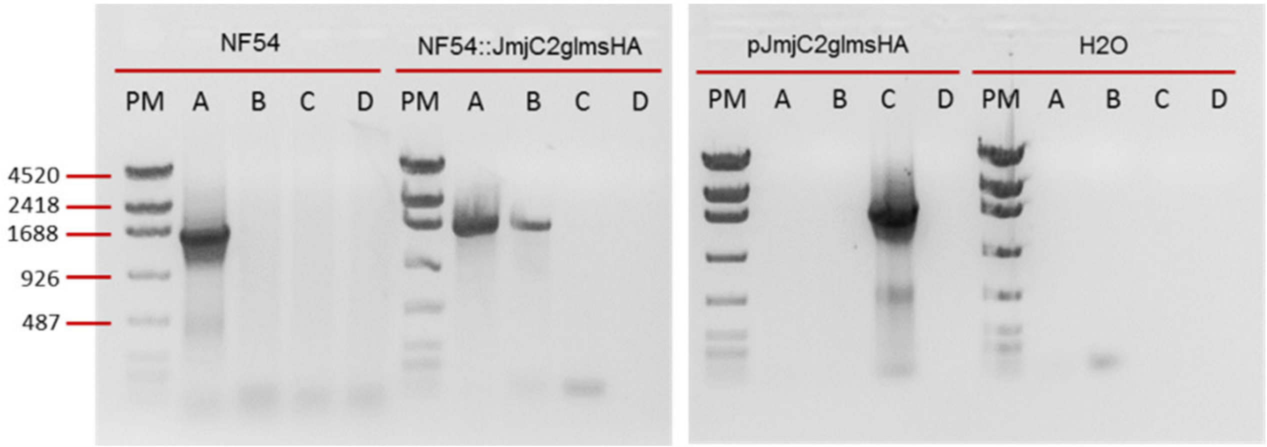


**A - primers 5 & 4 (1113 nt), B - primers 5 & 2 (1202 nt), C - primers 1 &2 (1267nt)**

C

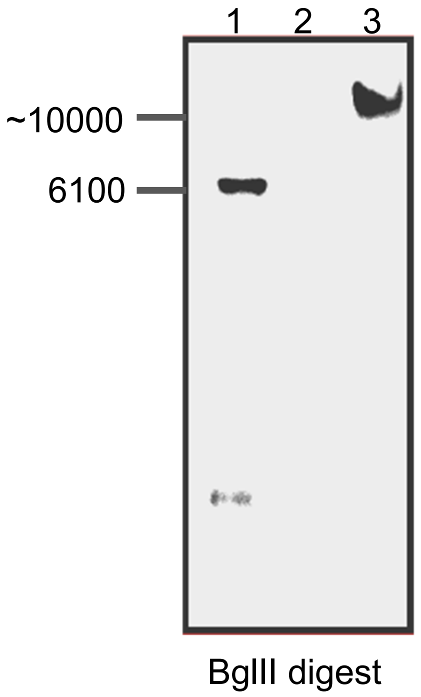


1 - plasmid DNA pJmjC2HAglmS
2 - gDNA WT NF54 control
3 - transfected gDNA of cloned transfectant NF54::JmjC2HAglmS

Supplementary Figure 1: Construction (**A**) and PCR analysis (**B**) of the tagged JmjC2 transfectant line. Oligonucleotides for PCR analysis are indicated. In **C,** Southern blot analysis using the DIG-labelled indicated probe on Bgl2 cleaved plasmid DNA (lane 1), genomic DNA of the NF54 line (lane 2) and lane 3, genomic DNA of the JmjC2HAglmS transgenic line.

### Supplementary Figure S2

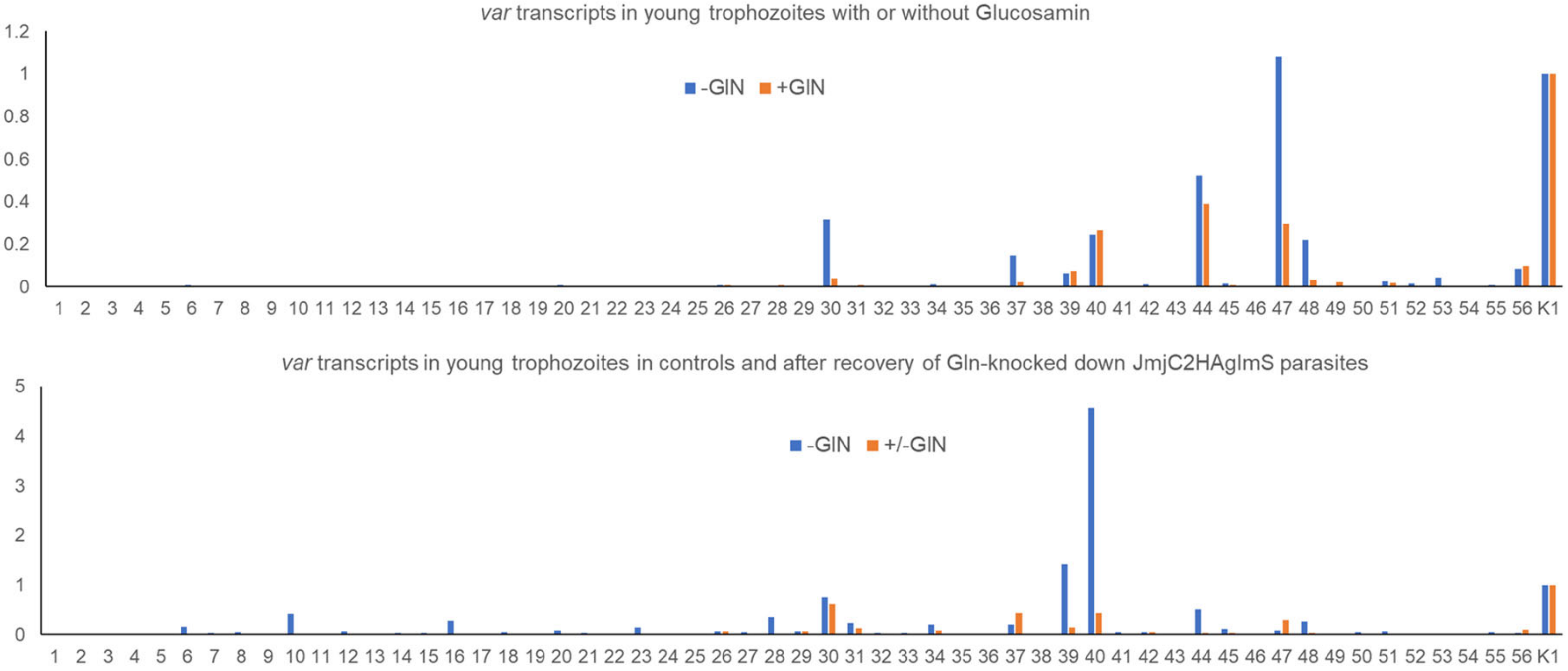


Supplementary Figure 2: *var* transcript profile with parasites outgrown for 15 cycles after cytoadherence selection over CHO-CD36 cells. RT-PCR analysis was done as described in Methods. K1 is the endogenous control transcript (*P. falciparum* t-seryl synthetase). Note that in this experiment, 10 reinvasion cycles occurred between parasites treated or not with Glucosamine. Also, note that the y-scale differs between the upper and the lower graph.

Supplementary Table 1: This table shows the real time PCR oligos used for var transcription analysis. These were originally created by Salanti and colleagues (2003, cited in the main text).

| NoLocu | former varID | Gene ID | fwdOligo | revOligo |
|---|---|---|---|---|
| 01 | PFA0015c | PF3D7_0100300 | TCATTATGGGAAGCACGATT | TGATTTCTACCATCGCAAGG |
| 02 | PFI1820w | PF3D7_0937600 | TGACCAAGACGAAGTATGGAA | TTGATCTCTGTTCGCTGTCC |
| 03 | PFD0020c | PF3D7_0400400 | ATATGGGAAGGGATGCTCTG | TGAACCATCGAAGGAATTGA |
| 04 | PFD1235w | PF3D7_0425800 | AAACACGTTGAATGGCGATA | GACGCCGAGGAGGTAAATAG |
| 05 | PFE1640w | PF3D7_0533100 | AAGAAAGTGCCACAACATGC | GTTCGTACGCCTGTCGTTTA |
| 06 | PF08_0141 | PF3D7_0800200 | GGTGTCAAGGCAGCTAATGA | TATGTCCTGCGCTATTTTGC |
| 07 | PF11_0008 | PF3D7_1100200 | GACGGCTACCACAGAGACAA | CGTCATCATCGTCTTCGTTT |
| 08 | PF11_0521 | PF3D7_1150400 | TGCTGAAGACCAAATTGAGC | TTGTTGTGGTGGTTGTTGTG |
| 09 | PF13_0003 | PF3D7_1300300 | CACAGGTATGGGAAGCAATG | CCATACAGCCGTGACTGTTC |
| 10 | PFA0005w | PF3D7_0100100 | TGCGCTGATAACTCACAACA | AGGGGTTCATCGTCATCTTC |
| 11 | PFA0765c | PF3D7_0115700 | AACCCCCAATACCATTACGA | TTCCCCACTCATGTAACCAA |
| 12 | PFB0010w | PF3D7_0200100 | ATGTGCGCTACAAGAAGCTG | TTGATCTCCCCATTCAGTCA |
| 13 | PFB1055c | PF3D7_0223500 | CAATTTTGGGTGTGGAATCA | CACTGGCCACCAAGTGTATC |
| 14 | PFC1120c | PF3D7_0324900 | CAATCTGCGGCAATAGAGAC | CCACTGTTGAGGGGTTTTCT |
| 15 | PFD0005w | PF3D7_0400100 | GACGACGATGAAGACGAAGA | AGATCTCCGCATTTCCAATC |
| 16 | PFD1005c | PF3D7_0421100 | ACCAAGTGGTGACAAAGCAG | GGGTGGCACACAAACACTAC |
| 17 | PFD1245c | PF3D7_0426000 | TGACGACTCCTCAGACGAAG | CTCCACTGACGGATCTGTTG |
| 18 | PFE0005w | PF3D7_0500100 | GAAGCTGGTGGTACTGACGA | TATTTTCCCACCAGGAGGAG |
| 19 | Mal6P1.316 | PF3D7_0600200 | TGGAAAGAACATGGACCTGA | TTCCTCGAGGGAAGAATCAC |
| 20 | Mal6P1.4 | PF3D7_0632500 | ATGTGTGCGAGAAGGTGAAG | TGCCTTCTAGGTGGCATACA |
| 21 | Mal6P1.1 | PF3D7_0632800 | GACAAATACGGCGACTACGA | TGTTTCACCCCATTCTTCAA |
| 22 | Mal7P1.50 | PF3D7_0712300 | ACCAAATGGTGACTTGCTCA | TTTTCATCGACGGATGATGT |
| 23 | Mal7P1.55 | PF3D7_0712800 | ACGTGGTGGAGACGTAAACA | CCTTTGTTGTTGCCACTTTG |
| 24 | Pf07_0050 | PF3D7_0712400 | GCGACGCTCAAAAACATTTA | TCATCCAACGCAATCTTTGT |
| 25 | PF07_0051 | PF3D7_0712600 | CGTGGTAGTGAAGCACCATC | CCCACCTTCTTGTGGTTTCT |
| 26 | PF08_0103 | PF3D7_0809100 | TGCAAGGGTGCTAATGGTAA | CCTGCATTTTGACATTCGTC |
| 27 | PF08_0106 | PF3D7_0808700 | TTTGTCCGGAAGACGATACA | ATCTGGGGCAGAATTACCAC |
| 28 | PF08_0140 | PF3D7_0800300 | TTTGGGATGACACCAAGAAA | GTCGCTTGATGAAGGAGTCA |
| 29 | PF08_0142 | PF3D7_0800100 | GTCGTGGAAAAACGAAAGGT | TATCTATCCAGGGCCCAAAG |
| 30 | PFI0005w | PF3D7_0900100 | TGCAAACCACCAGAAGAAAG | GTTCTCCGTGTTGTCCTCCT |
| 31 | PFI1830c | PF3D7_0937800 | CACACGTGGACCTCAAGAAC | AAAACCGATGCCAATACTCC |
| 32 | PF10_0001 | PF3D7_1000100 | GACGAGGAGTCGGAAAAGAC | TGGACAGGCTTGTTTGAGAG |
| 33 | PF10_0406 | PF3D7_1041300 | GTGCACCAAAAGAAGCTCAA | ACAAAACTCCTCTGCCCATT |
| 34 | PF11_0007 | PF3D7_1100100 | GAGGCTTATGGGAAACCAGA | AGGCAGTCTTTGGCATCTTT |
| 35 | PFL0005w | PF3D7_1200100 | CGGAGGAGGAAAAACAAGAG | TGCCGTATTTGAGACCACAT |
| 36 | PFL0020w | PF3D7_1200400 | TCGATTATGTGCCGCAGTAT | TTCCCGTACAATCGTATCCA |
| 37 | PFL0935c | PF3D7_1219300 | GACGCCTGCACTCTCAAATA | TTGGAGAGCACCACCATTTA |
| 38 | PFL1950w | PF3D7_1240300 | AGCAAAATCCGAAGCAGAAT | CCCACAGATCTTTTCCTCGT |
| 39 | PFL1960w | PF3D7_1240600 | CATCCATTACGCAGGATACG | AAATAGGGTGGGCGTAACAC |
| 40 | PFL1955w | PF3D7_1240400 | AAAGCCACTAGCGAGGGTAA | TGTTTTTGCCCACTCCTGTA |
| 41 | PFL2665c | PF3D7_1255200 | GGCACGAAGTTTTGCAGATA | TTTGTGCGTCTTTCTTCGTC |
| 42 | PF13_0001 | PF3D7_1300100 | ACAAAGGAACGTCCATCTCC | GCCAATACTCCACATGATCG |
| 43 | PF13_0364 | PF3D7_1373500 | CGGAATTAGTTGCCTTCACA | CATTGGCCACCAAGTGTATC |

| | | | | |
|---|---|---|---|---|
| 44 | PFD0615c | PF3D7_0412400 | ACCGCCCCATCTAGTGATAG | CACTTGGTGATGTGGTGTCA |
| 45 | PFD0625c | PF3D7_0412700 | TAAAAGACGCCAACAGATGC | TCATCGTCTTCGTCTTCGTC |
| 46 | PFD0630c | PF3D7_0412900 | ACTTTCTGGTGGGGAATCAG | TTCACCGCCACTTACTTCAG |
| 47 | PfD0995c | PF3D7_0420700 | TCACAACCTGACCCCCTACT | TCTTCGTCGTTGTCATCCTC |
| 48 | PFD1000c | PF3D7_0420900 | AGAGGGTTATGGGAATGCAG | GCATTCTTTGGCAATTCCTT |
| 49 | PFD1015c | PF3D7_0421300 | TGCAACGAAACATTAGCACA | AGCAGGGGATGATGCTTTAC |
| 50 | Mal6P1.252 | PF3D7_0617400 | ATTTGTCGCACATGAAGGAA | AACTTCGTGCCAATGCTGTA |
| 51 | Mal7P1.56 | PF3D7_0712900 | CACACATGTCCACCACAAGA | ACCCTTCTGTGGTGTCTTCC |
| 52 | PF07_0049 | PF3D7_0712000 | GTTGAGTCTGCGGCAATAGA | CTGGGGTTTGTTCAACACTG |
| 53 | PF07_0048 | PF3D7_0711700 | CAATTTTTCCGACGCTTGTA | CACATATAGCGCCGTCCTTA |
| 54 | PF07_0051 | PF3D7_0712600 | CGTGGTAGTGAAGCACCATC | CCCACCTTCTTGTGGTTTCT |
| 56 | PFL0030c | PF3D7_1200600 | TGGTGATGGTACTGCTGGAT | TTTATTTTCGGCAGCATTTG |
| K1 | PF07_0073 | PF3D7_0717700 | AAGTAGCAGGTCATCGTGGTT | TTCGGCACATTCTTCCATAA |
